# Preclinical Evaluation of Chemoradiation Resistance Using 18F-FDG PET/CT in Head and Neck Squamous Cell Carcinoma

**DOI:** 10.64898/2026.08.03.742521

**Authors:** Casey C. Heirman, Ashlyn G. Rickard, Rico Castillo, Katherine Gonzalez, Allison Pittman, Justin Smith, Katelyn E. Kelly, Jack B. Stevens, Tammara Watts, Yvonne M. Mowery, Kyle J. Lafata

## Abstract

**Purpose:** There is an urgent need for improved prognostic tools and biological understanding of chemoradiation resistance in head and neck squamous cell carcinoma (HNSCC). This study established a preclinical imaging dataset aimed at identifying prognostic imaging features from ^18^F-FDG micro-PET/CT scans in HNSCC mouse models.

**Methods:** Three orthotopic murine models were utilized: two human papillomavirus (HPV)-negative (MOC1, MOC2) and one HPV-positive (MLM1). When tumor volume exceeded 50 mm³, chemoradiation was initiated using cisplatin (5 mg/kg) and image-guided radiation therapy (8 Gy) on days 0 and 7. ^18^F-FDG micro-PET/CT imaging was performed on day 14. Tumors were manually segmented on PET/CT, and quantitative image features including tumor volume, SUVmean, and SUVmax were extracted. Treatment response was evaluated by relative tumor size on day 11 compared to day 0. Tumor growth and survival were compared across models using multiple-effects model and log-rank. Imaging feature associations were evaluated by Mann–Whitney U tests. Associations between survival, SUVmax, and tumor volume were assessed using Cox proportional hazards modeling and Kaplan–Meier analysis with log-rank testing.

**Results:** A total of 121 mice were treated and imaged. Significant differences in tumor growth and survival were observed among the three models (p < 0.01 for pairwise growth comparisons; p < 0.0001 for survival). Day 11 treatment response groups demonstrated significantly different growth trajectories following chemoradiation (p < 0.0001). SUVmax was significantly associated with survival (p = 0.0009), whereas SUVmean was not significant (p = 0.13). PET tumor volume demonstrated the strongest association with survival (p < 0.0001). A multivariate Cox proportional hazards model incorporating SUVmax and tumor volume significantly stratified survival risk (p < 0.0001).

**Conclusion:** Overall, these findings demonstrate that ^18^F-FDG PET/CT-derived metrics, particularly SUVmax and tumor volume, are robust predictors of chemoradiation response in orthotopic murine models of HNSCC.

## 1. Introduction

A major challenge in the treatment of head and neck squamous cell carcinoma (HNSCC) is resistance to chemoradiation therapy. Standard treatment typically includes surgery and/or scisplatin-based chemoradiation, with the goal of achieving long-term locoregional control while minimizing toxicity. Unfortunately, many patients do not respond to treatment or develop recurrence after initial complete response, indicating intrinsic or acquired therapeutic resistance [1]. Tumor hypoxia represents one key driver of resistance and poor therapeutic response in HNSCC [2, 3, 4]. In addition, human papillomavirus (HPV) status strongly influences clinical outcomes, with HPV-negative tumors demonstrating lower response rates and worse prognosis compared with HPV-positive oropharyngeal cancer [4, 5]. At the same time, intensified treatment strategies are associated with significant toxicities and reduced quality of life [6, 7], emphasizing the need for improved predictive biomarkers and more personalized therapeutic approaches. Emerging technologies, such as advanced imaging and computational tumor phenotyping, may help address this challenge by improving treatment response prediction and promoting better characterization of HNSCC tumor biology [8, 9].

Among imaging biomarkers, standardized uptake value (SUV), derived from ^18^F-fluoro-2-deoxy-glucose (FDG) positron emission tomography (PET), serves as a surrogate marker of tumor metabolism, which is particularly relevant in HNSCC [10, 11]. Elevated maximum SUV (SUVmax) has been linked to worse prognosis, suggesting metabolically active tumors are more treatment-resistant [12, 13]. In addition, mean SUV (SUVmean) values have been shown to provide potential prognostic value in HNSCC [12, 14, 15].

In this study, we address limitations of clinical studies by leveraging preclinical orthotopic models of HNSCC, in which tumor biology, PET/CT imaging, treatment conditions, and response dynamics are experimentally controlled. By integrating longitudinal tumor growth measurements and quantitative image analysis, we investigate the prognostic significance of PET/CT across three biologically distinct tumor models. We hypothesized that this approach would identify biomarkers that stratify therapeutic response across heterogeneous tumor biology in HNSCC.

## 2. Methods

**Figure 1.**
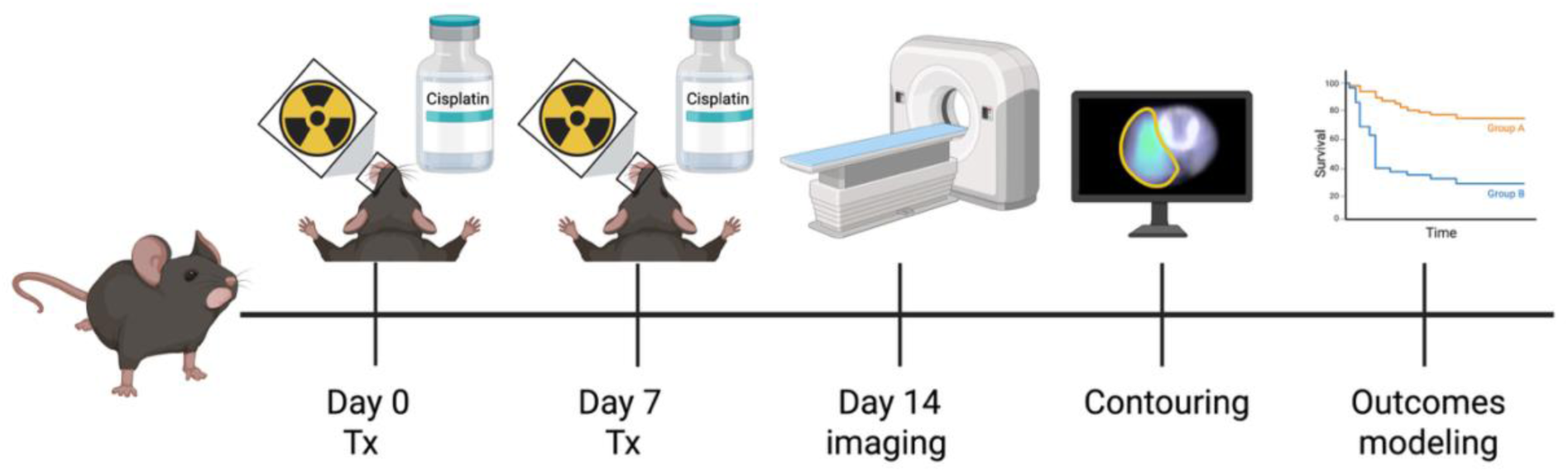
Experimental design to investigate chemoradiation resistance on PET/CT in preclinical tumor models of HNSCC. Mice bearing orthotopic tumors underwent chemoradiation treatments of 8 Gy radiation and 5mg/kg cisplatin on Days 0 and 7, followed by PET/CT imaging on Day 14. Subsequently, the tumor was contoured on PET/CT images, features extracted, and outcomes modeled and analyzed [16].

### 2.1 Tumor Induction and Monitoring

Mouse oral carcinoma (MOC) cell lines, MOC1 and MOC2, were sourced from Kerafast, while the MLM1 (mEERL lung metastasis clone 1) cell line was generously provided by Dr. Paola Vermeer at Sanford University. MOC1, developed in C57BL/6 mice through carcinogen exposure, demonstrates an indolent growth pattern characterized by increased infiltration of CD8+ T-cells into the tumor microenvironment, inducible MHC class I expression, and reduced CD4+ T-cell presence [17]. In contrast, MOC2 exhibits aggressive behavior, including decreased MHC class I expression, infiltration of FOXP3+CD4+ regulatory T-cells, and the presence of CD11b+/Gr1+ cells, indicative of immunotolerance [17]. MLM1 is an HPV-positive and immune-competent cell line expressing HPV16 E6 and E7 proteins [18]. These proteins contribute to P53 degradation, affecting cell cycle regulation and tumor suppression mechanisms [18].

MOC cells were maintained at 37°C with 5% CO_2_ in media consisting of 500mL Iscove’s Modified Dulbecco’s Medium (Gibco) supplemented with 250mL F-12 Nutrient Mix (Gibco), 37.5mL Fetal Bovine Serum (FBS), 7.5mL penicillin/streptomycin (P/S, Gibco), 0.33mg/mL hydrocortisone (Sigma-Aldrich), 10mg/mL insulin (Sigma-Aldrich), and 0.2 mg/mL human EGF (Gibco). MLM1 cells were maintained at 37°C with 5% CO_2_ in media consisting 507mL Dulbecco’s Modified Eagle’s Medium (Gibco) supplemented with 168mL Hams F-12 Nutrient Mix (Gibco), 75mL FBS, 7.5mL P/S, 0.33mg/mL hydrocortisone, 25mg/mL transferrin, 10mg/mL insulin, 0.2mg/mL tri-iodo-thyronine, and 0.2mg/mL human EGF. All animal studies were approved by the Duke University Institutional Animal Care and Use Committee (IACUC) and adhered to the NIH Guide for the Care and Use of Laboratory Animals. Viable MOC1 (300,000 cells/50μL/mouse in 1:1 PBS:Matrigel [Corning]), MLM1 (300,000 cells/50μL/mouse in 1:1 PBS:matrigel), or MOC2 (30,000 cells/50μL/mouse in PBS) were submucosally injected into the right buccal mucosa of male and female C57BL/6J mice (Jackson Laboratories, strain #000664), aged between 6 to 10 weeks, while anesthetized with <3.5% isoflurane in 100% oxygen carrier gas. Following induction, mice were monitored thrice weekly for tumor development. Tumor dimensions were measured using digital calipers, and volume was approximated using equation 1.

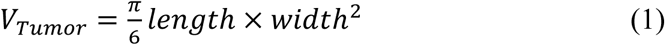

Mice were euthanized upon tumor ulceration, significant weight loss (>20%), or tumors exceeding 12 mm in any dimension.

### 2.2 Treatment and Imaging

Mice received two fractions of 8 Gy radiotherapy administered one week apart on Days 0 and 7 with concurrent cisplatin (5 mg/kg, intraperitoneal) administered 30 minutes prior to irradiation. Mice were irradiated with opposed lateral beams (4 Gy/field) using the Xstrahl Small Animal Radiation Research Platform (SARRP) at 220kV and 13mA, while anesthetized with 2% isoflurane. Prior to each irradiation, setup was confirmed via on-board fluoroscopy. Daily saline boluses of 0.5mL were delivered subcutaneously for two days following cisplatin injections to minimize nephrotoxicity.

^18^F-FDG PET/CT imaging was performed on Day 14 post-treatment. Mice were fasted overnight, injected with ^18^F-FDG via tail vein, and imaged using a microPET/CT scanner (Siemens Inveon) under isoflurane anesthesia.

### 2.3 Quantitative Image Analysis

Initial regions of interest (ROIs) were segmented based on asymmetry visible on micro-CT imaging. Within the initial ROIs, Hounsfield unit (HU) thresholding was employed, selecting only voxels within the ROI with HU values falling between 75-300, and smoothed with a gaussian filter with a standard deviation of 2.0, creating the final ROI used in analysis. These HU thresholds were selected based on experimental data and literature review [19, 20, 21, 22].

Cylindrical ROIs were also segmented in the liver for subsequent SUV normalization to each individual mouse’s metabolic baseline [23]. Raw PET counts were converted to SUV. SUV metrics from tumor ROIs were then normalized to SUVmean for liver ROIs to generate a final metabolic intensity map, from which SUVmax, SUVmean, and tumor volume were extracted.

### 2.4 Statistical Analysis

Statistical analyses were performed using MATLAB (MathWorks) and GraphPad Prism. Image analyses utilized Gremse-IT Imalytics Preclinical software. Associations between PET metrics (SUVmax, SUVmean), tumor volume, and tumor model (MOC1, MOC2, MLM1) were evaluated. Tumor growth and survival were compared across models using mixed-effects model and log-rank, and imaging feature associations were evaluated using Mann–Whitney U tests.

Treatment response and survival outcomes were assessed using Kaplan–Meier analysis with log-rank tests and Cox proportional hazards regression. Statistical significance was defined as *P* < 0.05, with *P*-values adjusted for multiple comparisons using Bonferroni corrections where appropriate.

## 3. Results

### 3.1 Radiation Response of Different Mouse Models

Experimental data were collected for 121 mice (MOC1, *n* = 43; MOC2, *n* = 38; MLM1, *n* = 40) that underwent treatment and imaging. Two MOC1 mice were excluded from downstream image analysis because their tumors were cured prior to imaging on day 14. The mean time from tumor induction to treatment was 15.7 ± 10.0 days, with a median of 12 days. Among the three tumor models, MOC1 exhibited the longest induction period (18.5 ± 11.7 days), followed by MOC2 (15.0 ± 8.8 days) and MLM1 (11.0 ± 5.5 days).

Following treatment initiation, median time to endpoint was 35 days for the entire cohort, with MOC1 exhibiting the longest survival time at a mean of 54.2 ± 34.2 days, compared with 30.7 ± 7.5 days, for MOC2 and 31.8 ± 8.8 days for MLM1 (Table 1). The time from treatment to endpoint in MOC1 was ∼41–43% longer than that of MOC2 and MLM1 and also demonstrated greater variability, whereas MOC2 and MLM1 showed more consistent survival outcomes.

**Table 1.**
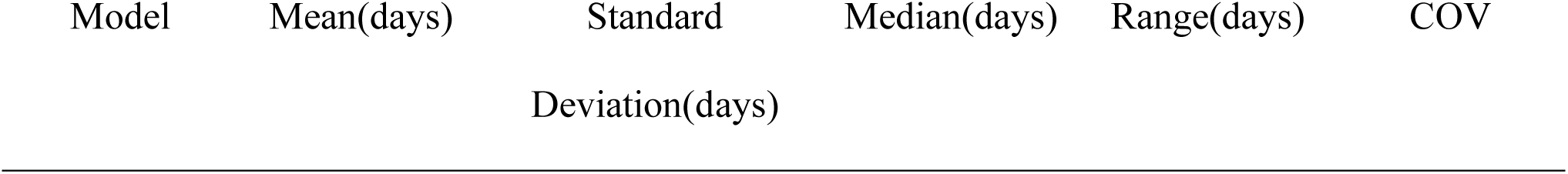
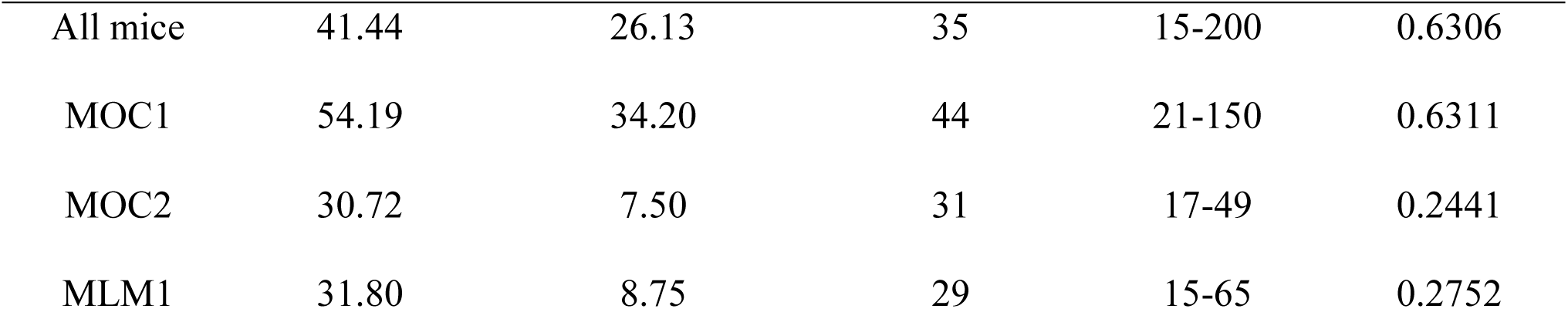
Post-treatment survival.

| Model | Mean(days) | Standard<br>Deviation(days) | Median(days) | Range(days) | COV |
| --- | --- | --- | --- | --- | --- |
| All mice | 41.44 | 26.13 | 35 | 15-200 | 0.6306 |
| MOC1 | 54.19 | 34.20 | 44 | 21-150 | 0.6311 |
| MOC2 | 30.72 | 7.50 | 31 | 17-49 | 0.2441 |
| MLM1 | 31.80 | 8.75 | 29 | 15-65 | 0.2752 |

Significant differences in growth patterns were identified among the three models (MOC1 vs. MOC2, MLM1 vs. MOC1 and MOC1 vs. MOC2, p ≤ 0.0001; mixed-effects model, Figure 2A). Significant differences using mixed-effects model and Tukey’s post hoc analysis revealed all three models to be significantly different by day 11. Using day 11 normalized growth, “High responders” demonstrated no external asymmetry after treatment, whereas “mid responders” exhibited tumor shrinkage, and “low responders” showed continued tumor growth 11 days after treatment initiation. Significant growth trend differences were observed across these three groups (high and mid responders p ≤ 0.0001, mid and low responders p ≤ 0.0001, high and low responders p ≤0.001, mixed-effects model, Figure 2B). Additionally, the three models had significantly different survival (p ≤ 0.0001, log-rank), with significant differences for MLM1 vs. MOC1 (p ≤ 0.001), MLM1 vs. MOC2 (p ≤ 0.001), and MOC1 vs. MOC2 (p ≤ 0.0001), Figure 2C). Median survival was 31 days for MOC2, 38 days for MLM1, and 43 days for MOC1.

**Figure 2.**
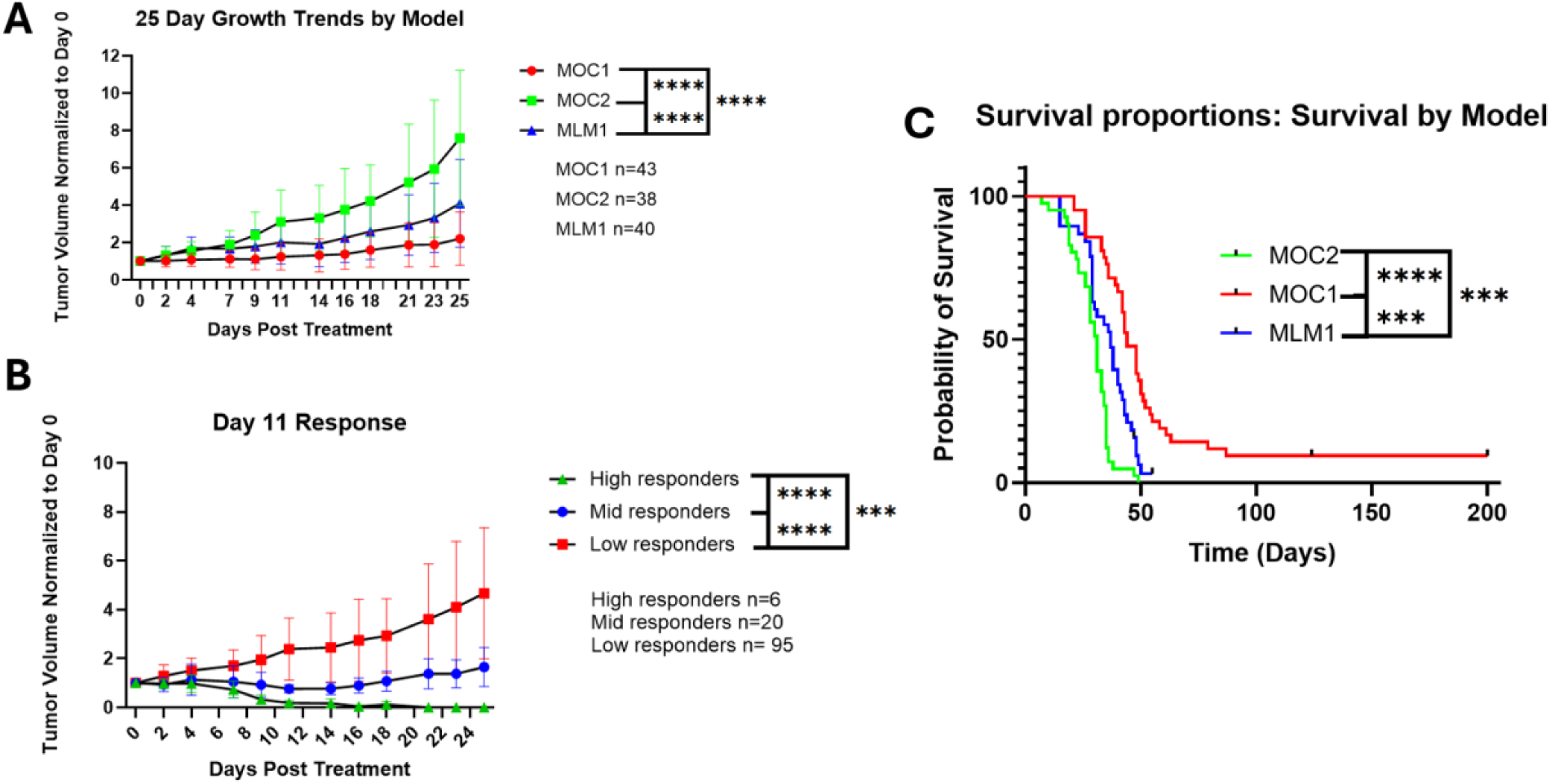
Tumor responses of the different immunological phenotypes. *A.* Tumor growth trends by tumor model, error bars represent the standard deviation. *B.* Tumor growth by early response, which is defined as tumor volume change on day 1**1** compared to the normalized volume on the day of initiation. *C.* Mouse survival after treatment initiation by tumor model.

### 3.2 Imaging Feature Analyses

Descriptive statistics for the extracted PET/CT imaging features are summarized in Table 2. The average tumor volume on Day 14 was 111 mm³ (median: 94 mm³), with values ranging from 12 to 401 mm³. Mean tumor SUVmax and SUVmean were 4.13 and 2.17, respectively, while liver SUVmax and SUVmean demonstrated lower variability, with mean values of 0.81 and 0.57, respectively. The mean normalized tumor SUVmax, calculated relative to liver uptake, was 7.67 (range 2.56 to 31.74).

**Table 2.** Descriptive statistics for image features extracted from the PET/CT regions of interest in the tumor and liver for the complete dataset.

|  | **Volume** | **Tumor** | **Tumor** | **Liver** | **Liver** | **Normalized** |
| --- | --- | --- | --- | --- | --- | --- |
|  | **(mm<sup>3</sup>)** | **SUVmax** | **SUVmean** | **SUVmax** | **SUVmean** | **SUVmax** |
| **Mean** | 110.83 | 4.13 | 2.17 | 0.81 | 0.57 | 7.67 |
| **Median** | 93.71 | 3.70 | 2.09 | 0.82 | 0.58 | 6.68 |
| **Minimum** | 11.53 | 0.14 | 0.07 | 0.05 | 0.04 | 2.56 |
| **Maximum** | 400.51 | 10.50 | 4.24 | 1.42 | 1.10 | 31.74 |

SUVmax (p = 0.0009, log-rank, Bonferroni-corrected; Figure 3A) was a significant predictor of survival when placed in high risk and low risk groups using the mean risk score, whereas SUVmean (Figure 3B) was not significantly associated with survival (p = 0.13, log-rank, Bonferroni-corrected). Tumor volume was also a significant predictor of survival (p = 6.54e-20, log-rank, Bonferroni-corrected, Figure 3C). The multivariate Cox proportional hazards analysis (Figure 3D) demonstrated that both SUVmax and tumor volume were associated with risk of death, and Kaplan–Meier survival analysis using median risk score as the stratification threshold showed clear separation between survival for the high-risk and low-risk groups (p = 4.44e-20, log-rank, Bonferroni-corrected).

**Figure 3.**
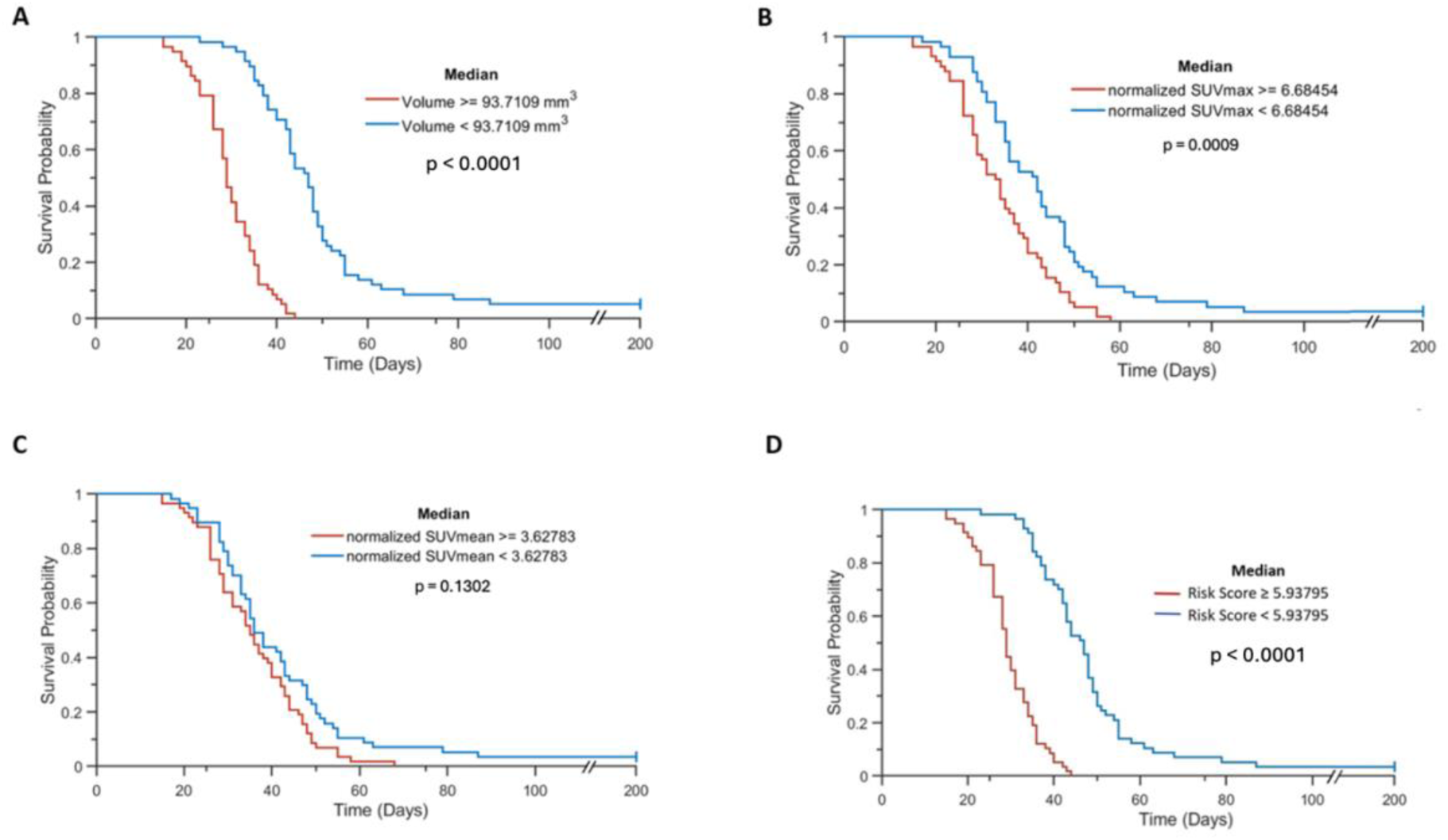
Survival based on first order intensity and shape features. *A.* Survival based on median tumor volume on CT scan from Day 14. *B*. Survival for groups based on median normalized tumor SUVmax value on PET scan from Day 14. *C.* Survival based on median tumor SUVmean on PET scan from Day 14. *D.* Survival based on Cox proportional hazard model median risk score using SUVmax and tumor volume on Day 14 as covariates. Both higher tumor SUVmax and tumor volume at Day 14 were associated worse survival. SUVmax demonstrated a hazard ratio of 1.0538 (Cox coefficient = 0.0524), indicating that increasing metabolic activity was associated with a higher risk. Similarly, increasing tumor volume was associated with increased worse survival (hazard ratio = 1.0151; Cox coefficient = 0.015), suggesting that larger tumor burden at this early time point may be associated with poorer outcomes.

Both higher tumor SUVmax and tumor volume at Day 14 were associated worse survival. SUVmax demonstrated a hazard ratio of 1.0538 (Cox coefficient = 0.0524), indicating that increasing metabolic activity was associated with a higher risk. Similarly, increasing tumor volume was associated with increased worse survival (hazard ratio = 1.0151; Cox coefficient = 0.015), suggesting that larger tumor burden at this early time point may be associated with poorer outcomes.

Only the MOC1 and MOC2 models showed significant differences in tumor volume by CT on Day 14, with MLM1 falling in between the two MOC models (Figure 4A). No significant differences were observed in SUVmax across models (Figure 4B).

**Figure 4.**
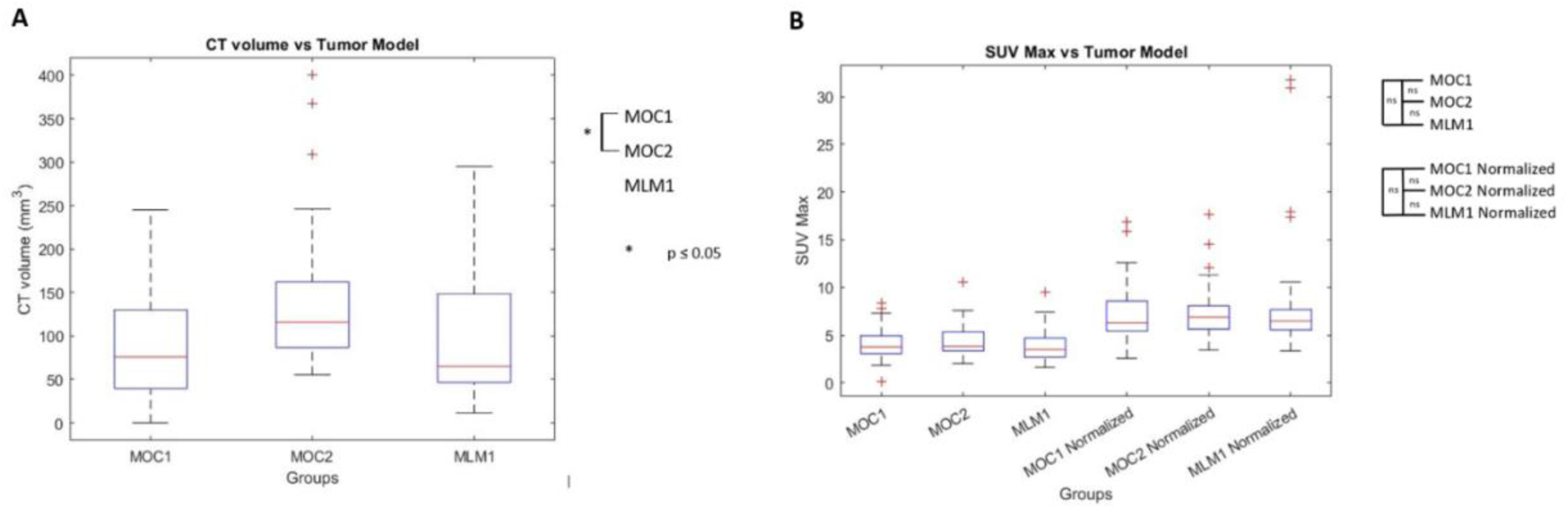
***A.*** Tumor volume based on ROI on the CT scan on Day 14 across tumor models. ***B.*** SUVmax (with and without normalization to liver SUVmean) vs tumor model.

## 4. Discussion

^18^F-FDG PET/CT-derived imaging biomarkers play an increasingly important role in the characterization, monitoring, and assessment of treatment response in HNSCC [24]. However, interpretation of these imaging biomarkers remains challenging due to the biological heterogeneity of HNSCC and dynamic nature of tumor response during treatment [25]. In this work, we developed a preclinical framework for evaluation of ^18^F-FDG PET/CT imaging biomarkers in controlled, syngeneic, orthotopic mouse models of HNSCC with variable therapeutic responses. Our approach enabled assessment of post-treatment tumor metabolism and tumor growth over time, while minimizing confounding variability commonly present in clinical patient populations, such as heterogeneous imaging protocols, limited longitudinal data, and comorbidities. We demonstrated the feasibility of this approach by evaluating the relationship between SUVmax, SUVmean, tumor volume, and overall survival following chemoradiation.

MOC1 mice exhibited significantly longer median survival and delayed tumor growth pattern compared to MOC2 mice, which is consistent with prior studies demonstrating the more aggressive and less immunogenic phenotype of MOC2 tumors relative to MOC1 tumors [17, 26]. MLM1 tumors exhibited intermediate growth kinetics (Fig 2A) and survival relative to MOC1 and MOC2 (Fig 2C), which is consistent with prior characterization of MLM1 as a distinct immunological phenotype with intermediate tumor aggressiveness [18]. Additionally, each mouse model exhibited intra-model heterogeneity regarding growth rates. This variability may more closely reflect the biological heterogeneity observed clinically in HNSCC, where differences in tumor metabolism, immune microenvironment, and intrinsic sensitivity to (chemo)radiation contribute to variable treatment outcomes [27].

Tumor volumetry has demonstrated prognostic value in HNSCC, with larger baseline tumor volumes associated with poorer outcomes [28, 29, 30]. Longitudinal assessment of tumor growth kinetics further improves response characterization by capturing dynamic changes in tumor burden rather than static measurements alone [31, 32, 33, 34]. Accordingly, preclinical head and neck tumor models commonly rely on longitudinal volumetric growth as a measure of treatment efficacy [35, 36, 37].

Prior radiobiological studies have demonstrated that measurable divergence in tumor growth kinetics can emerge within days of irradiation in preclinical tumor models [38]. Similarly, murine studies using subcutaneous UT-SCC-14 tumors treated with cisplatin-based chemoradiation reported early treatment-induced growth delay, with clear separation of tumor growth curves from untreated controls by day 11 after treatment initiation [39]. Clinical imaging studies have also observed significant tumor volume reductions approximately two weeks after treatment initiation [40]. Together, these findings support the use of early volumetric changes as meaningful indicators of treatment response that may precede long-term survival outcomes. Consistent with these observations, CT-derived tumor volume at day 14 was a strong predictor of overall survival in our study.

Beyond volumetric assessment, metabolic imaging also provided prognostic information. Higher post-treatment SUVmax values were associated with faster tumor growth and reduced overall survival, consistent with clinical studies demonstrating that increased FDG uptake reflects more aggressive tumor biology and poorer prognosis in HNSCC [41]. In contrast, SUVmean was not significantly associated with overall survival. Because SUVmean is averaged across the entire tumor ROI, metabolically inactive regions such as necrosis may dilute its value and reduce its prognostic utility [42]. These findings suggest that localized regions of elevated metabolic activity, rather than average uptake alone, better reflect biologically relevant tumor heterogeneity, consistent with previous studies highlighting the prognostic significance of metabolic heterogeneity in HNSCC [8, 9].

From a mechanistic perspective, previous studies have reported significant correlations between higher FDG uptake and alterations in genes such as TP53, as well as positive associations between FDG uptake and genetic heterogeneity [43]. Alterations in metabolic genes have also been associated with changes in the tumor immune microenvironment, with high-risk HNSCC profiles demonstrating reduced infiltration by several immune cell populations, including CD8+ T cells, mast cells, and NK cells [44]. Additionally, specific T-cell profiles within tumor-draining lymph nodes have been associated with differences in tumor size [45]. Together, these findings suggest that metabolic imaging biomarkers such as FDG uptake may reflect broader genomic and immunologic features associated with tumor aggressiveness and progression. One possible explanation for the differences observed among tumor models is the distinct molecular profile of the HPV-positive MLM1 model, which lacks functional p53 [18]. As a tumor suppressor, p53 plays a central role in regulating cellular metabolism by promoting oxidative phosphorylation and suppressing glycolysis, the primary metabolic pathway utilized by many cancer cells [46]. Consequently, disruption of this pathway may contribute to differences in FDG uptake and treatment response, although this relationship was not directly evaluated in the present study.

Although this study makes significant strides in preclinical evaluation of HNSCC, several limitations should be considered when interpreting the findings. First, the study was conducted in orthotopic transplant mouse models. While advantageous for enabling controlled experimental conditions and longitudinal monitoring, transplant models do not fully recapitulate the complexity and heterogeneity of human HNSCC. Factors that influence treatment response in patients, including gradual tumor development under immune surveillance, are therefore not fully represented in this system. By contrast, genetically engineered mouse models, in which oncogenes or tumor suppressor genes are selectively altered, or carcinogen-induced models may better capture the genetic and biological heterogeneity of human HNSCC, while still allowing control over factors such as comorbidities and treatment regimen [47, 48]. Second, pretreatment imaging was not performed in this study due to logistical limitations. Previous studies have reported conflicting findings regarding the prognostic value of pretreatment PET imaging in HNSCC, with some analyses demonstrating limited prognostic utility [8, 49], while others have identified significant associations between pretreatment SUV parameters and clinical outcomes [14, 50]. Evaluation of pretreatment imaging within this controlled experimental environment is highly valuable to stratify baseline metabolic activity in relation to treatment response.

Normalizing post-treatment SUV measurements to the tumor’s baseline could improve our ability to predict treatment response and survival.

## 5. Conclusion

Utilizing small animal imaging systems and preclinical tumor models provided an experimental framework to assess imaging biomarkers of chemoradiation response and resistance. This approach has potential to enhance our understanding of the metabolic and volumetric indicators that underly chemoradiation resistance in HNSCC.

